# Interior soft x-ray tomography with sparse global sampling

**DOI:** 10.1101/2025.08.11.669333

**Authors:** Axel Ekman, Jian-Hua Chen, Carolyn A. Larabell, Mark A. Le Gros, Venera Weinhardt

**Affiliations:** National Center for X-ray Tomography, Lawrence Berkeley National Laboratory, Berkeley, CA, USA; Department of Anatomy, University of California, San Francisco, San Francisco, CA, USA; Centre for Organismal Studies, Heidelberg University, 69221 Heidelberg, Germany

## Abstract

Objective. To investigate the feasibility of interior imaging reconstruction in soft X-ray tomography to achieve higher spatial resolution cellular imaging, including whole-cell imaging. Approach. We develop an alignment and reconstruction algorithm that enables a combination of a low number of images from sparse whole-cell imaging with a high-resolution local interior scan. Based on numerical simulations, we demonstrate that combined reconstructions mitigate the depth of field limitation in high-resolution scans, enable radiation dose optimization, and yield quantitative X-ray absorption values with sparse sampling. Furthermore, we validate our numerical approach using experimental data from two different cell types and demonstrate that combined reconstruction is a reliable method for obtaining high and local spatial resolution within the volume of a whole cell. Significance. The developed sparse reconstruction algorithm provides a robust and faithful visualization of cellular organelles with soft X-ray tomography. A mesoscale imaging approach, such as an interior tomography scan, enables one to “scout” and zoom into the volumes of interest that contain organelles of interest. Utilizing sparse reconstructions, this increase in spatial resolution is achieved without sacrificing larger volume imaging, providing information on the relative position of all organelles within a cell.

## 1 Introduction

Over the past years, soft X-ray tomography, or SXT, has established itself as a powerful imaging technology to tackle key questions in cell biology. To date, SXT has been used on more than 100 different cell types^45^. Several features set SXT apart from other imaging modalities. The so-called “water-window” energy range used in SXT enables native contrast of cellular anatomy without the need for labeling or chemical fixation. This native contrast is quantitative and is extensively used to study the state of cellular organelles^16,17^ and mathematical modeling of their molecular composition^1^. Furthermore, SXT employs transmission geometry and computed tomography (CT) acquisition to visualize whole cells with tens of nanometers spatial resolution.

This similarity to medical CT comes along with comparable limitations. The magnification and camera pixel size limit the spatial resolution of SXT. Due to the use of diffractive X-ray optics in SXT, the increase in spatial resolution comes not only with a smaller field of view on the camera sensor but also with a shorter depth of focus. Therefore, whole-cell SXT imaging with higher spatial resolution can be achieved only on smaller specimens, like bacterial cells^46^. The loss of imaging volume in high-resolution SXT can be compensated by alternative imaging geometries that combine several volumes imaged in depth^33^ or by laterally expanding the field of view^11^. However, these approaches often result in higher radiation doses or loss of spatial resolution in some parts of the specimen. Therefore, the challenge of SXT imaging lies in increasing spatial resolution without loss of imaging volume or an increase in radiation dose, inevitably leading to incomplete data.

In medical CT, the incomplete data is often solved by alternatives to filtered-back-projection reconstruction algorithms, such as optimal recovery, Bayes estimate, and Tikhonov-Phillips methods^29^. In a specific case where high-resolution CT scans are acquired locally, known as region of interest (ROI) tomography, the out-of-field structures affect the quantitative accuracy of the X-ray absorption coefficient and lead to artifacts, particularly at the edges of the field of view^15^. Kyrieleis et al. ^22^ show that a simple extension of the truncated data can be sufficient for high-quality reconstructions using standard reconstruction methods. This approach, however, calls for a 4x increase in the number of projection images acquired to obtain accurate X-ray absorption values, which is often not feasible for SXT due to the increase in radiation dose.

For correct quantitative reconstruction, multi-resolution approaches where an interior ROI CT scan is supplemented by the scout scan of a whole sample at lower resolution were proposed^3,8^. Several interior reconstruction methods were designed to utilize the multi-resolution data. Using the low-resolution data as a prior in the reconstruction of the ROI scan^25^ or re-projecting sparse views to get extended data for ROI scan^40^ helps to reduce artifacts from data truncation and provide high-quality, reliable reconstructions of interior tomography at low computational costs.

Interestingly, the dose-fractionation theorem that is valid for biological specimens measured in computed tomography geometry^18,27^ suggests that the dose required to reconstruct a high-resolution 3D volume can be distributed among any number of different projections. Thus, an accurate reconstruction of X-ray absorption values is possible without an increase in radiation dose for such multi-resolution approaches.

Despite the broad applicability of ROI tomography in medical and laboratory CT imaging and the possibility of combining projections at no increase in radiation dose, this imaging approach has not been employed in SXT. On the one side, flat specimen supports used in some SXT instruments do not allow for full profit from multi-resolution imaging, as the samples are laterally extended. On the other hand, full-rotation SXT imaging at higher resolution is limited not only by the short depth of field but also by the mechanical stability of the microscope^46^.

Here, we develop and optimize the reconstruction algorithm, combining sparse low-resolution and interior high-resolution SXT scans to achieve accurate and stable interior tomography in SXT. Based on theoretical considerations, we find an optimal number of low-resolution images required to obtain high-fidelity, high-resolution local imaging. Furthermore, we show that dose distribution optimization enables multi-resolution interior SXT implementation.

Finally, we probe our theoretical considerations experimentally by performing interior SXT tomography of bacteria and human B cells. We show that our algorithm provides a theoretically exact interior SXT reconstruction that is reliable and has great potential for cell imaging where high and local spatial resolution is crucial, such as the substructure of small bacterial cells and membrane structures within larger human cells.

## 2. Method

In X-ray tomography, the image formation model has traditionally been based on the Radon transform^38^, the ideal linear transform (projection) of the specimen’s attenuation coefficients onto a plane. This is linked to the experimental image formation through the Beer-Lambert law, such that the recorded intensity of a ray, *I*_*i*_, can be expressed as attenuation of its intensity along a ray path, *L*_*i*_, as

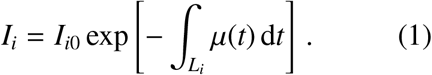

In this work, for the inversion, we consider only the measurement in terms of a linear transform on the discrete representation of the X-ray linear absorption coefficient (LAC) distribution ***x*** such that

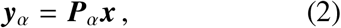

where the matrix elements *P*_*αij*_ represent the contribution of *j*th voxel in the LAC distribution on the projection on the *i*th measured pixel, and ***y***_*α*_ is a vector representation of the measured absorption image − log ***I***_*α*_*/****I***_*α*0_.

A tomographic measurement can now be expressed as a series of projection operators

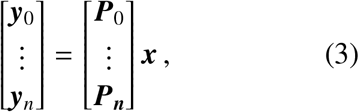

where ***P***_*n*_ is the measurement matrix describing the image formation of the *n*th image and ***y***_*n*_ its corresponding absorption image. The tomographic inversion is then described by its “measurement matrix”

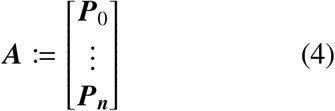

and thus, with a set of linear equations

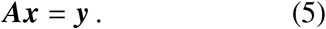

The behavior of ***A*** and the possible existence of its inverse depend on the measurement setup. In general, no unique solution exists for an overdetermined system because of noise, but suitable solutions can be found, e.g., via the normal equation

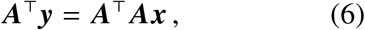

which describes the critical points of the *l*^2^-norm of the measurement errors.

The least-squares solution is not unique for an undetermined system, and the solution depends on the initial point and the reconstruction algorithm. Examples of such undetermined measurements, such as insufficient sampling, limited angle acquisition, and interior tomography, are shown in Fig. 1.

**Figure 1.**
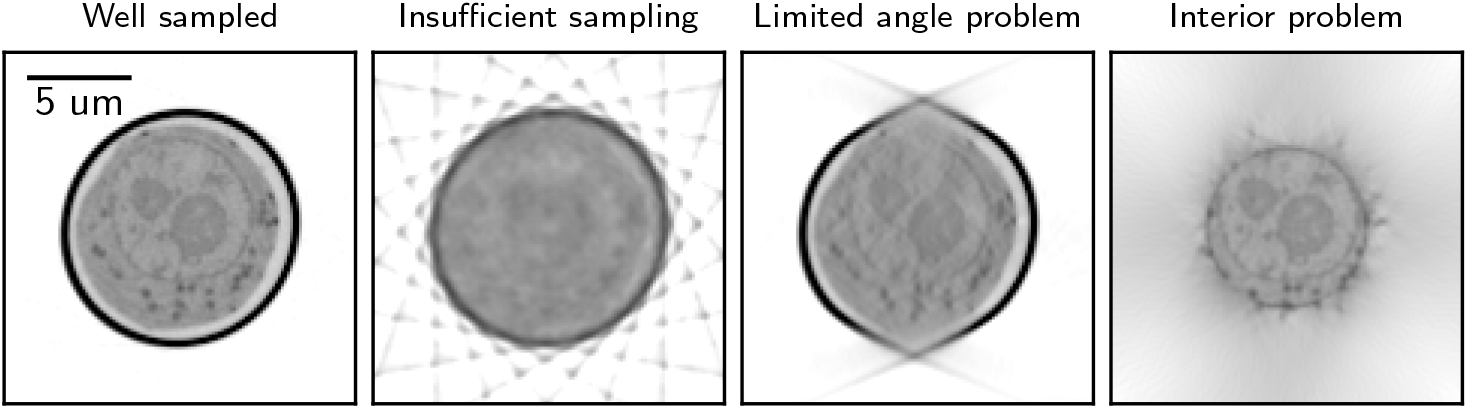
Examples of undetermined measurements in SXT, showing typical reconstruction artifacts arising from the nature of the forward model ***A***. From left to right: Sufficient well-sampled reconstruction with 92 projection images acquired; insufficient sampling, where the number of projections was reduced to 10; limited angle problem where 100 projections were acquired from − 65^°^ to 65^°^, and Interior problem where the detector width is reduced to 8 µm (from 16 µm).

In X-ray tomography, the interior problem is nearly solvable, with the primary challenge being a low-frequency ‘cupping’ bias^26^, which can be mitigated using lambda tomography^13^ or by incorporating known X-ray attenuation in subregions^7^.

In this study, we explore whether incorporating a sparse full field-of-view (FOV) scan can provide sufficient stabilization for the reconstruction process, particularly in high-resolution quantitative soft X-ray tomography experiments.

Specifically, we do this by extending the measurement matrix ***A***_ROI_ containing the interior scans with a set of full FOV scans ***A***_full_, such that

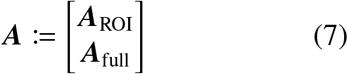

Our work here focuses primarily on the feasibility of the inverse problem in this setting, the sampling considerations for the SXT case, and an experimental proof of concept. Thus, all reconstructions are performed using a pure *l*^2^ minimization of the measurement matrix via the conjugate gradient method on the normal equations (CGNE). For the simulated results, we determine the optimal stopping iterations based on the *l*^2^ loss.

While numerous reconstruction approaches incorporate, e.g., more accurate statistical modeling^14^, regularization^50^, or deep learning^41^, our focus remains on the fundamental tomography model as a straightforward linear inverse problem. This formulation aims to provide a generalizable framework that can readily adapt to more advanced iterative schemes.

### 2.1. Null space

To assess the stability of the reconstruction within the region of interest (ROI), we evaluate its contribution to the null space of the projection operator ***A***. As discussed in^21^, the domain 𝕌 of the measurement matrix ***A*** can be divided into two subspaces, its null space *N*(***A***) and its measurable space *N*_⊥_(***A***), where the null space is formally defined as

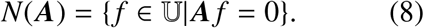

This implies that for any vector ***x*** ∈ 𝕌 that is a solution of Eq. (5), then{***x*** + ***x***_null_} is also a solution to the equation for any vector ***x***_null_ ∈ *N*(***A***). Therefore, any solution component residing in the null space does not affect the projection data and thus cannot be recovered from the measurements alone.

To estimate the null space, we follow the methods discussed in^49^ and^21^ by initializing the ***x***_0_ with a phantom image. We update the image using Wilson-Barrett iterations^48^

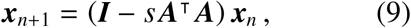

where *s* is an iteration step size. From the starting point of the phantom image ***x***_0_ = ***x***_meas_ + ***x***_null_, the system can only update measurable components of ***x***_0_, thus converges towards the null space component ***x***_null_ of the phantom.

We conducted simulations with bandwidth-limited, noiseless projection images, reconstructing them using CGNE. The number of projections was selected based on the sampling requirements for a bandwidth-limited 2D Radon transform^39^. The reconstruction process was halted at the point of highest peak signal-to-noise ratio (PSNR). In Fig. 2, we show both the PSNR and the relative length of the null space vector, |***x***_*null*_| */* |***x***_0_| for different SXT measurement setups: a full-view scan ***A***_full_, an interior scan ***A***_ROI_, and a combination of sparse full-view and interior scans as defined in Eq. (7).

**Figure 2.**
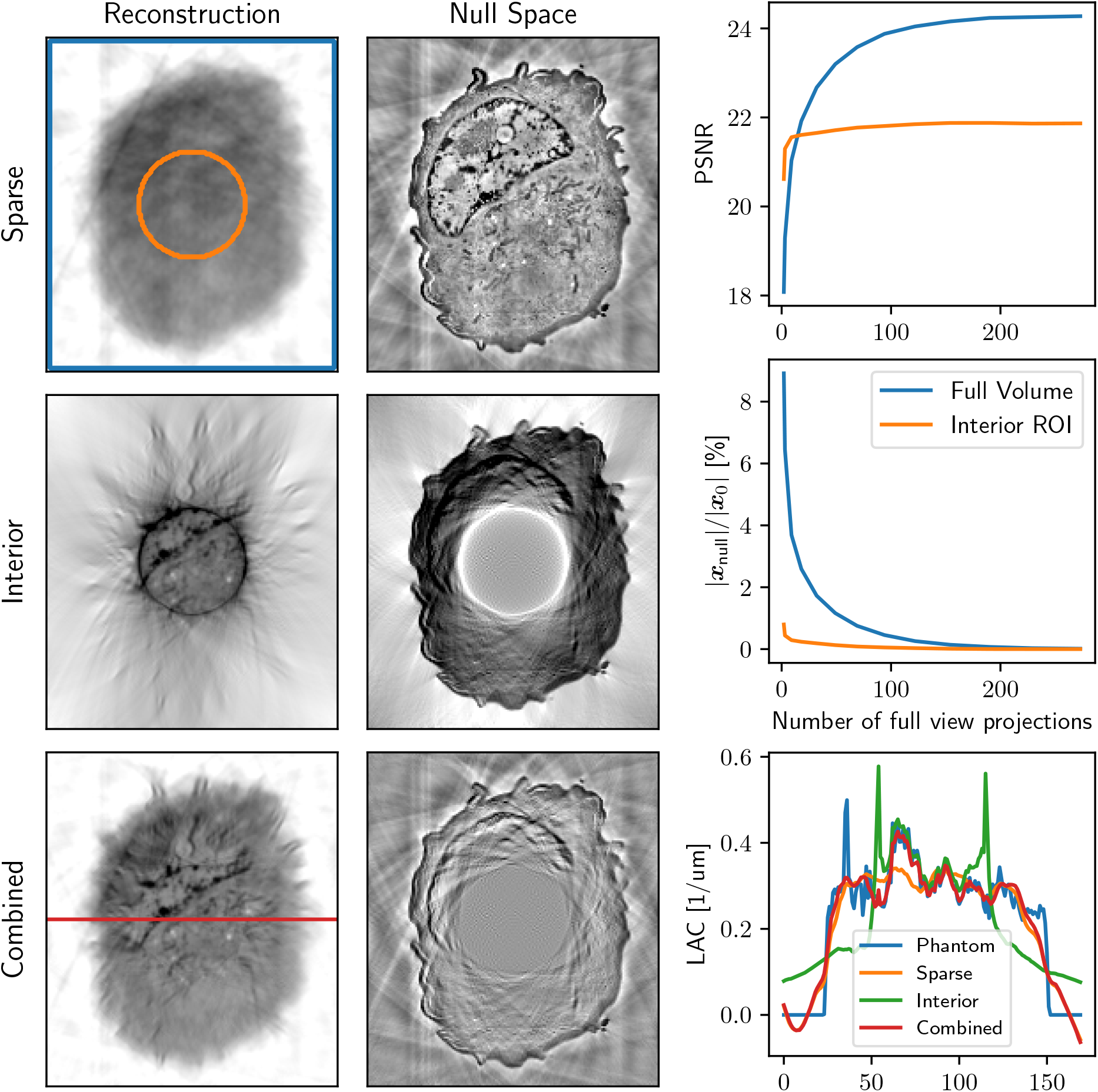
Left: Examples of reconstructions and their corresponding null space components, ***x***_null_, for a sparse low-resolution scan (7 projection images), an interior scan of the same phantom (273 truncated projection images), and a combined reconstruction of sparse low resolution and interior tomography (7 full and 273 truncated images). Right top: PSNR of the ROI within the interior scan (orange circle) and the area outside the ROI (blue square), both for the combined reconstruction as a function of the number of projections used in the (sparse) full FOV scan. Right, middle: the relative length of the null space vector for the inside and outside ROI (orange circle). Right, bottom: LAC profile (across the horizontal red line) for the original phantom and three SXT reconstructions, which are sparse low-resolution imaging, interior high-resolution, and combined reconstructions.

From both the reconstructions and the profiles on Fig. 2, we observe that the sparse scan provides a reconstruction without bias but lacks detailed resolution. While having enough sampling to capture the fine details in the truncated data, the interior scan exhibits the common artifacts associated with interior tomography. However, by combining both measurement matrices, the artifacts are corrected, and the details of a sample are preserved in reconstruction.

Furthermore, we can see that although the PSNR (and, inversely, the length of the null space vector) converges gradually when measured across the full volume, the same metrics as measured from the interior ROI stabilize rapidly with just a few full-view scans. Notably, while the matrix itself is still underdetermined, remarkably few full FOV projections are required to achieve a stable reconstruction within the ROI of the interior scan. Importantly, the LAC values of combined interior SXT reconstructions are the closest to the selected phantom.

### 2.2 PSF considerations

X-ray microscopic images are not ideal projections of objects because they are influenced by the microscope’s three-dimensional point spread function (PSF)^32^. The resolution of the optical system, such as SXT, is determined by the relationship *r* ∝ *λ/NA*, where *λ* is the wavelength of the illuminating light and *NA* is the numerical aperture of the objective lens. However, diffraction-limited optics impose a maximum depth of field (DOF) described by *DOF* ∝ *λ/NA*^2^. This narrow DOF restricts the volume in which image formation can approximate parallel projections, and particularly for large samples, may introduce radial reconstruction artifacts^10^.

To address the impact of limited DOF on the accuracy of interior SXT tomography, we numerically evaluated the X-ray optics of the XM-2 microscope. The image formation process was modeled as an incoherent intensity transform, incorporating a PSF with added Poisson noise, following the approach in^10^. We analyzed two X-ray objective lenses with outer zone widths (OZW) of 35 nm and 60 nm, as described in Sec. (A).

To evaluate the quality of a reconstruction, ***x***, we define a relative peak signal-to-noise ratio PSNR_rel_ as

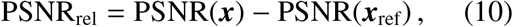

where ***x***_ref_ is the “ideal” reference reconstruction. For ***x***_ref_, we used a high-resolution that is independent of depth (infinite DOF) and has the same lateral resolution as the 35 nm OZW. Fig. 3 presents the numerical results of Eq. (10) for an 6 µm wide interior, showing variations in PSNR_rel_ as a function of the radius of the cylindrical measurement region.

**Figure 3.**
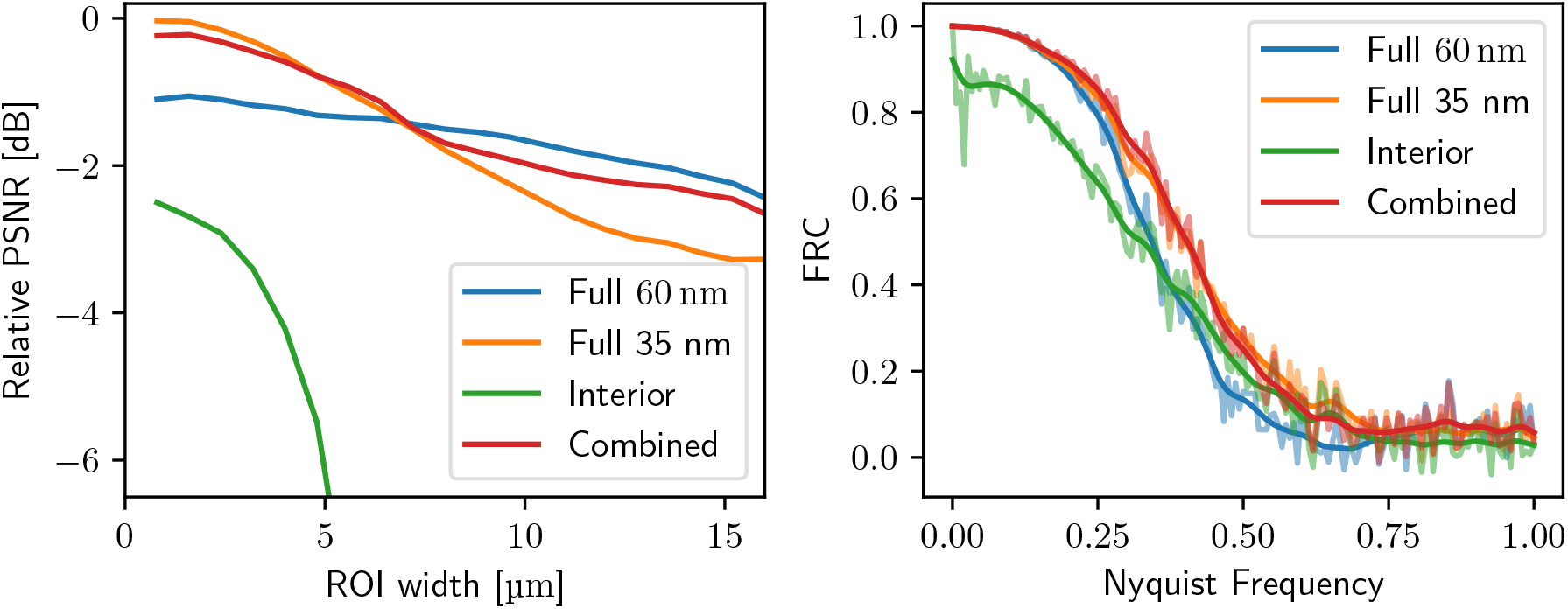
Left: PSNR comparison for different scanning modes in an interior tomography setup with a 6 µm ROI. The interior scan behaves as expected. The full scan with 35 nm OZW provides better interior detail but suffers from reduced quality because of the shorter DOF. The combined scan with truncated 35 nm data and full 60 nm data preserves high inner detail without artifacts of the interior scan. Right: Corresponding in-slice FRC with an ROI of 6 µm. The square ROI includes the interior boundary to exaggerate artifacts. The low-frequency bias explains the huge drop in PSNR. The multi-view protocol, including both 60 nm and 35 nm data, preserves both PSNR and FRC.

As expected, the PSNR and Fourier ring correction analysis (FRC) for the interior scan (in green) suffer due to bias from the truncated data, with image quality rapidly deteriorating towards the edges of the ROI. The full scan with a 35 nm OZW (in orange) scan provides better detail within the central region compared to the 60 nm OZW (in blue), but the resolution as measured by PSNR diminishes as a result of the shorter DOF.

The combined scan, using the truncated 35 nm data alongside the full 60 nm data, successfully captures the inner details of the high-resolution scan without the loss of image quality typically associated with the interior scan. That shows that the constraints of limited DOF go away when the interior SXT tomography is reconstructed in a combined fashion with the ‘out of focus’ knowledge from the sparse scan. As shown in Fig. 3, the combined reconstruction of the low and high-resolution projection matrix can slightly degrade the quality of the interior scan as compared to the ideal high-resolution case. This is due to the non-optimal weight of the back-projected measurements. Using the true PSF of the microscope with its inversion^10^ would mitigate this, as the back projection weights the overlapping interior data appropriately. In practice, however, this is computationally costly (on the order of *N*^2^, where *N* is the width of the kernel) and suffers from slower convergence. Another solution is to apply a data-weighting scheme, such as in Cao et al. ^3^, to take into account the different weights of the measurement data.

We suggest approximating the projection operator as

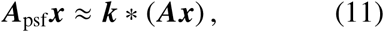

where the kernel ***k*** is a z-independent PSF and ∗ the 2D convolution operator. Conversely, the adjoint operator can be expressed as

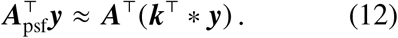

This 2D deconvolution approach drastically speeds up the projection operator and allows it to be more easily integrated into existing projection libraries.

### 2.3. Sampling considerations

The system’s optics set a physical limit on the achievable spatial resolution. However, in an experimental setup, this represents an optimistic upper limit. In reality, the spatial resolution is constrained by measurement statistics, influenced by the total radiation dose the sample can tolerate, and the specific sampling protocol used.

Dose fractionation tells us that we are free to distribute the dose in whichever way we want as long as we remain sufficiently sampled (for details, see Conf. Sec. (B.3)). However, experimentally, it is often beneficial to take fewer projections, as processing, such as alignment or deconvolution of the projection images, is much more robust. Fig. 12 shows FRC analysis of conjugate gradient applied to normal equation (CGNE) reconstruction method for different angular sampling and the radiation dose. Angular sampling shows no effect on FRC curves at a lower total dose.

### 2.4. Dose optimization

Achieving high-resolution imaging with full FOV scans may require higher doses than currently used. If one is only interested in the ROI of the scan, the dose distribution of a full scan is highly non-optimal. This is illustrated in Fig. 4 where we show the dose distribution across a sample for three different scanning approaches: half rotation with 180^°^ for the full FOV with 97 images scan, full rotation 360^°^ for the full FOV with 97 images, and a combined scan with 19 full FOV images, combined with 97 interior images. The intensity of the interior scans was increased so that the total absorbed dose in the sample was the same as for other scanning geometries.

**Figure 4.**
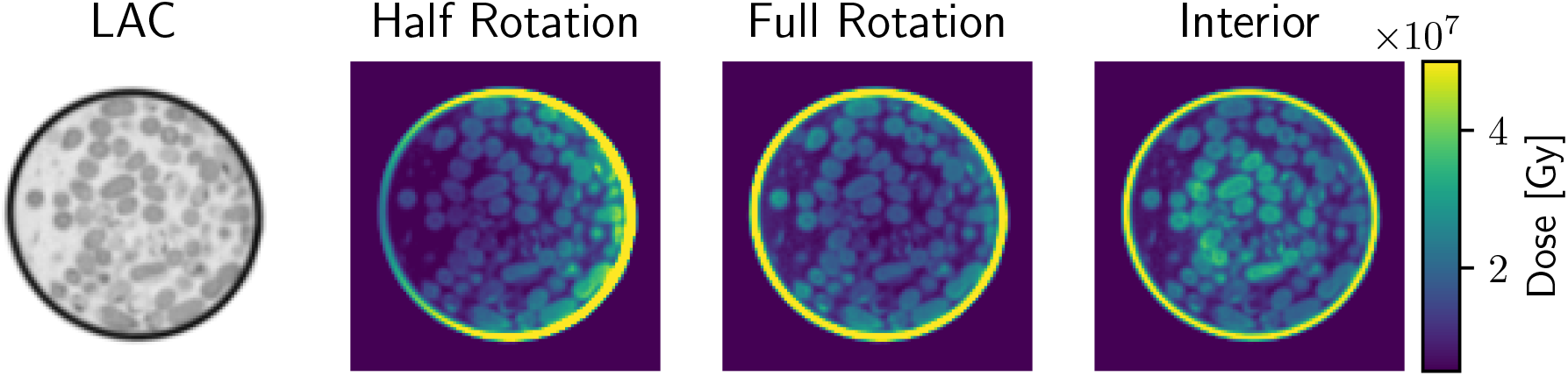
Comparison of the radiation dose distribution in a sample for different scanning protocols. On the left is simulated X-ray absorption (LAC) within a sample. On the right in color, there are three scanning geometries: half rotation (180 degrees), full rotation (360 degrees), and combined reconstructions. The total dose on the sample for all scanning geometries is the same. The full FOV scan concentrates the dose at the sample’s edges, leading to inefficient sampling. The interior scan distributes the dose more effectively within the center of the specimen.

It was previously reported^24^ that the highest dose levels accumulate at the sample edges during a half-rotation FOV scan. To optimize dose distribution, the most straightforward approach is to use a full 360-degree rotation protocol. In full rotation tomography, it is essential to interlace the mirrored images such that the angular sampling is increased (if mirrored images overlap, the angular sampling is halved). Despite this, even in full-rotation scanning, a significant portion of the radiation dose is still concentrated at the sample edges and specimen holder, increasing the risk of localized radiation damage.

Going one step further, the interior SXT scan distributes the dose more effectively across the specimen, focusing on areas where detailed imaging is required. Additionally, as discussed in Sec. (2.1), the quality of the ROI for the combined projection operator improves rapidly with just a few full FOV scans. Beyond this point, additional full-FOV sampling provides diminishing returns regarding ROI quality while unnecessarily increasing the total radiation dose.

To further investigate this trade-off, we simulated the accumulated dose in a highresolution phantom, comparing a full scan to an interior scan while keeping the incident intensity constant. For this phantom, the average accumulated dose of a full scan, *D*_*Full*_, was 2.3 times higher than that of the ROI scan, *D*_*ROI*_. This means that for each full FOV image removed, the remaining dose allows for a *D*_*Full*_*/D*_*ROI*_ increase in the intensity of the interior scan while keeping the total dose constant.

To maintain consistent angular sampling for the interior scan, the redistributed dose was evenly allocated among the interior projections, leading to the following relationship:

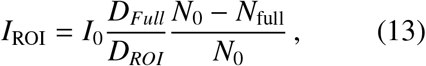

where *N*_0_ = 499 is the full angular sampling used as a reference.

In Fig. 5, we show the simulation phantom and the results of the dose optimization. As expected, while the PSNR of the full-FOV reconstruction (blue) increases with an increasing number of projection images, the quality of the combined interior scan (orange) is higher at the low number of projection images (**more than**) as the dose is distributed more efficiently (lower statistical error on the interior measurements). This beneficial trade breaks down at very low angular sampling (orange circle), as the number of full FOV projections is insufficient to reduce the interior bias.

**Figure 5.**
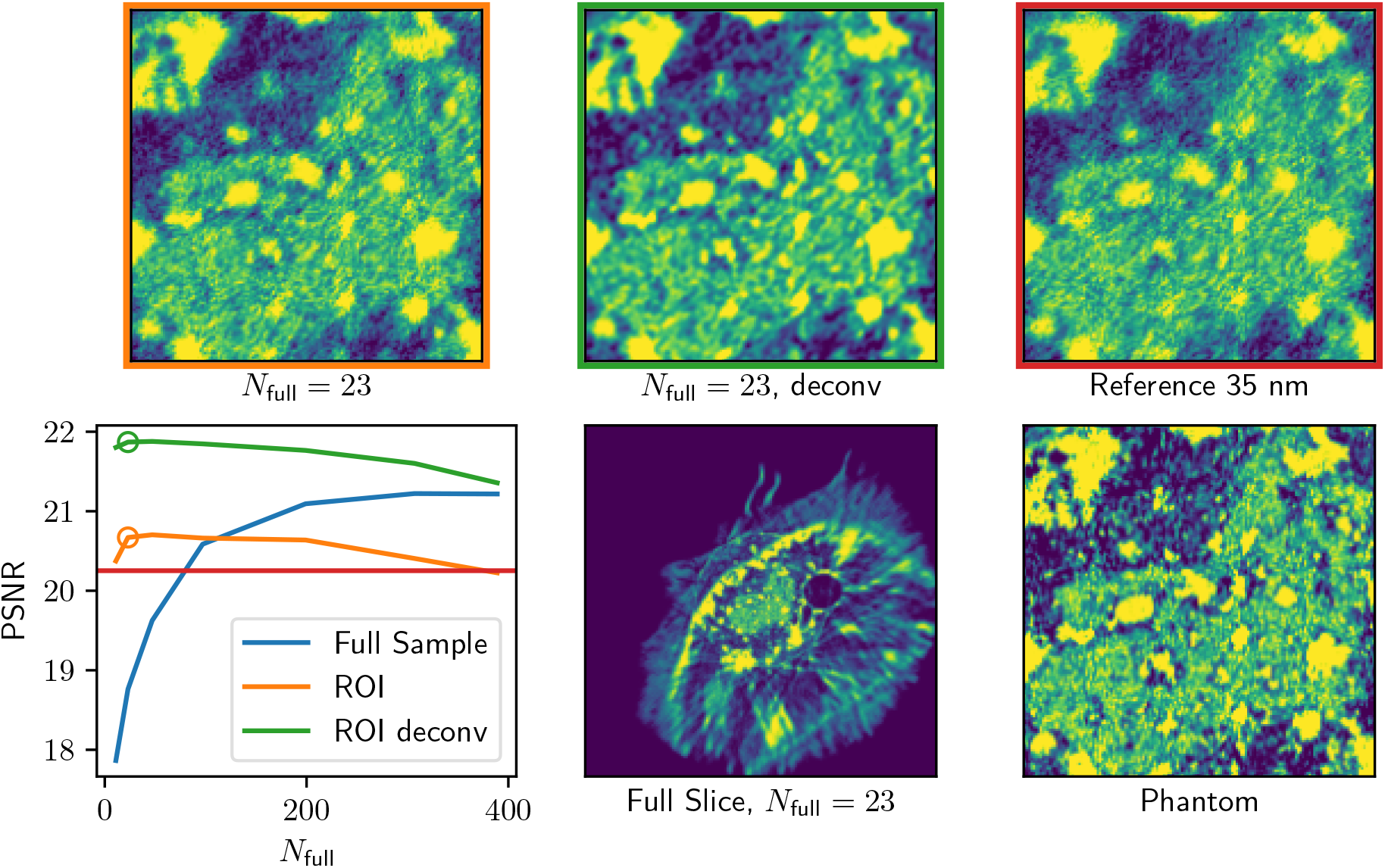
Comparison of full scan and dose-optimized scan performance. The protocols were designed so that the total dose of the sample was kept the same. The gained intensity from reducing the number of full scan images *N*_full_ was distributed evenly across the interior scans. From the PSNR curves, we see that the overall quality decreases as the angular sampling becomes sparse, but the interior PSNR increases. At some sparse sampling, this breaks down, as the sparse measurement matrix is insufficient to reduce the interior bias in the null space. The top row shows a detail of the ROI as compared to the reference scan, i.e., a full FOV 35 nm scan with infinite DOF.

Additionally, we show the result of the deconvolved reconstructions using 2D approximation as described in Eq. (12). To mitigate possible inversion crimes (where both the forward model and the backward model are done with the exact same discrete operator ***A***), the deconvolution reconstruction was done by using a Gaussian kernel with a full-width-half-maximum equal to OZW. This deconvolution approach significantly helps to improve the quality of the combined interior scan as measured using PSNR.

Overall dose and angular sampling optimization for the phantom measurements show that a combined interior scan can be achieved with only 23 full projections. In practice, the optimal selection of dose reduction is additionally limited by experimental factors, such as alignment, which is discussed in Sec. (B).

## 3. Experimental results

To verify the applicability of the interior SXT experimentally, we have applied our interior reconstruction method on bacteria (*Pseudomonas putida*, KT2440 strain) and human B lymphocytes (GM12878 from the NGIMS Human Genetics Cell Repository). These specimens demonstrate a variety of cells that would profit from interior SXT. That is, from small cells like bacteria and yeast, where a high number of tiny cells and their structure can be quantitatively analyzed, and larger human cells, where tiny structural changes appear in an unpredictable location within the larger cell volume, are visualized in the context of other organelles and a whole cell. Like other full-rotation SXT experiments, specimens were loaded into thin-wall glass capillaries and vitrified via rapid plunging into a liquid propane^5^.

For each approach, 92 projection images were acquired for full and interior tomography with 2^°^ rotation increment and 200 ms to 500 ms exposure time per projection. The projection images of full FOV and interior scans were aligned using a combination of an automatic alignment of the full FOV scans and cross-correlation of the interior projections to the full FOV data, described in detail in Sec. (B). To demonstrate the differences in interior SXT for each cell type, we reconstruct full FOV, interior SXT, and combined reconstruction based on a full interior scan and 19 projection images from full FOV.

Fig. 6 demonstrates the results obtained for an interior high-resolution scan, a sparse low-resolution SXT scan, and the combined interior tomography reconstruction. To enhance the demonstration of interior tomography, the ROI projections have been additionally cropped by 25% to reduce the FOV of the scans artificially. In-plane reconstructed virtual slices, as expected, show artifacts outside the imaging area in the interior scan. Although the interior reconstructions enable the visualization of cells inside the specimen holder, the addition of the sparse full FOV scan substantially increases the usable ROI of the reconstruction. It delivers faithful LAC reconstruction over the whole ROI.

**Figure 6.**
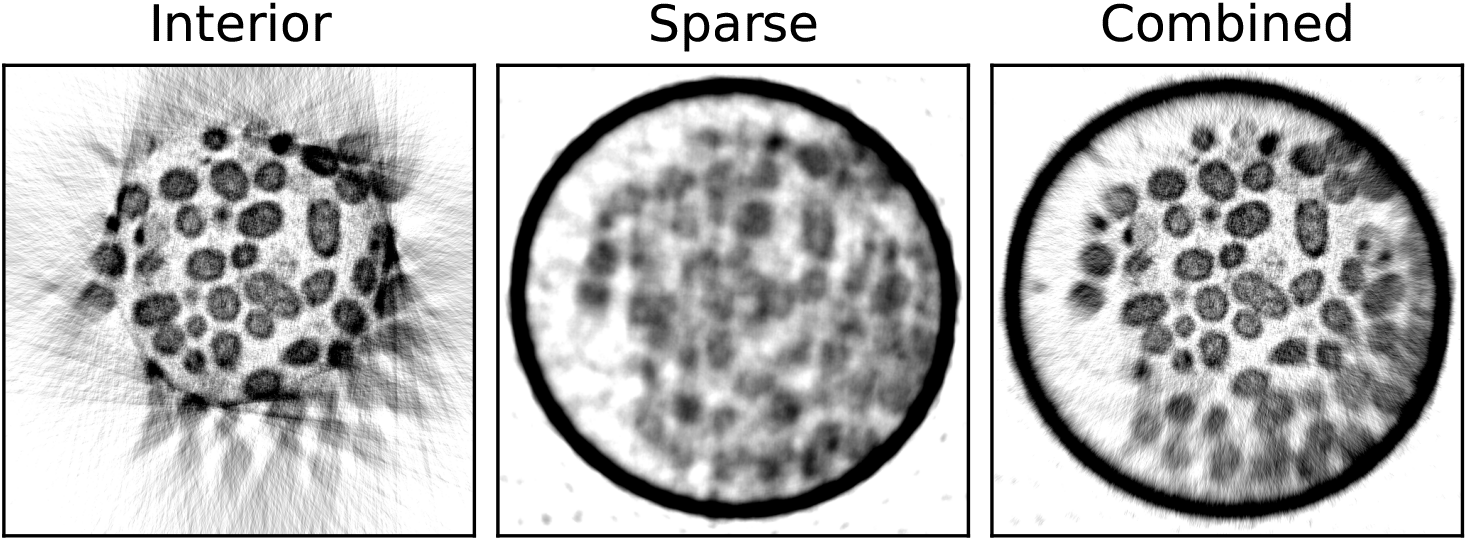
Interior SXT demonstrated experimentally on bacteria cells. Top: Transverse virtual slices through the SXT volume for interior and sparse scans only and combined reconstruction as described above. To emphasize the interior effects, the interior scan was reconstructed from 35 nm ROI images cropped horizontally by 25 %.

In one such interior SXT, there are tens of individual bacterial cells, each showing variable membrane aberrations. Such subcellular changes in bacteria are of high relevance to research on genetic mutants^6^, metabolism^12^, and bacterial biofilms^4^.

A similar comparison of the interior, sparse, and combined SXT imaging of a human B cell is shown in Fig. 7. All major organelles, such as the nucleus, mitochondria, and lipid droplets, are visible in all reconstruction examples. Smaller organelles, such as the endoplasmic reticulum (ER) and endosomes, are visible but challenging to segment and analyze quantitatively in sparse reconstruction. In comparison, when interior SXT is truncated with projections from full FOV, the reconstructed combined volume is free from artifacts with faithful LAC values and locally higher resolution.

**Figure 7.**
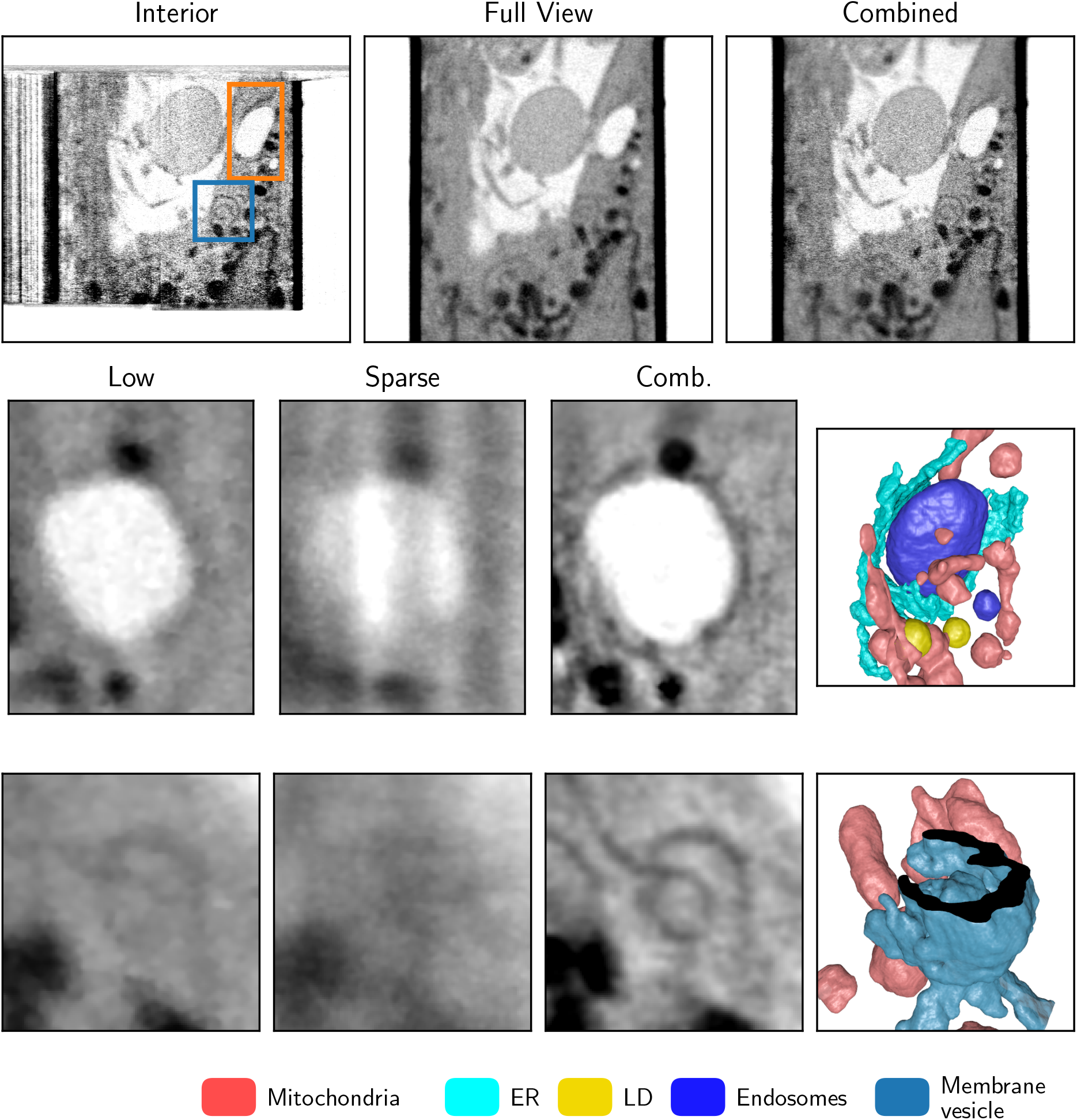
Interior SXT demonstrated experimentally on human B cells. Top: Saggital virtual slices through the SXT volume for interior and sparse scans only, combined reconstruction as described above, and two highlighted ROIs. The middle and bottom rows show a detailed ROI of ER wrapping around the late endosome (orange ROI) and a double membrane vesicle (blue roi) from low-resolution full FOV data, sparse global sampling, and combined interior reconstruction. Part of the outer membrane is clipped to reveal the structure inside.

This higher resolution region within the whole cell enables the segmentation of the region of interest (ROI1 and ROI2). Both ROIs show faint details in the low-resolution scan that could not be faithfully labeled but were segmentable semi-automatically in the combined interior scan with sparse global sampling. This type of interior high-resolution imaging enables the analysis of fine cellular features like membranes within whole human cells. Its combination with the sparse information enables a) to determine the region of interest for a “zoom in” high-resolution SXT scan; b) to retrieve contextual information, such as proximity to the nucleus or other organelles; and c) to reconstruct faithful LAC values, enabling automatic and quantitative analysis. For the example of the ER-endosome interaction network, even just the sparse scan is sufficient to probe the position of the desired ROI, as the position of the endosome does not require high resolution or angular sampling.

Overall, the experimental results show that the combined reconstruction developed for the interior SXT scans enables the visualization of subcellular features in a large contextual volume and can be robustly used for other scientific cases.

## 4. Conclusions

In this work, we have demonstrated via the Null space of the combined projection matrix that the bias in the interior tomography vanishes when combined with sparse context scans. We showed numerically that the limitation of the shallow depth of field in the highresolution interior scan is not relevant for the combined reconstructions, as the sparse scan provides the “out of focus” information. Furthermore, based on the dose fractionation theorem, we argue that since SXT imaging is primarily noise-limited, optimizing the dose distribution is more critical than increasing angular sampling. The calculated dose distribution confirms that the combined interior SXT reconstruction utilizes the radiation dose more effectively within the specimen holder than half-and full-rotation tomography.

In practice, the optimal selection of imaging is limited by experimental factors, such as alignment. We, therefore, performed multiresolution imaging of bacteria and human B cells. Based on these experimental data, we show that combined reconstructions of the interior SXT enable faithful reconstructions of LAC values in 3D. For small cells like bacteria or yeast, our combined reconstructions allow for analysis of subcellular alterations for tens of cells. Conversely, combined interior reconstructions required only 19 projections from the sparse scan. That enables us to perform low-resolution “scout” SXT imaging, followed by a zoom-in into the region of interest in larger cells, like human B cells. Therefore, the combined reconstruction of the interior SXT imaging is a valuable tool for several application cases.

Our combined reconstruction algorithm of interior tomography provides numerical consideration and the first experimental evidence that the resolution limit in SXT imaging can be increased without sacrificing larger-volume imaging.

## 5. Acknowledgment

Soft X-ray tomography was conducted at the National Center for X-ray Tomography, which is supported by NIH NIGMS (grant no. P30GM138441) and the Department of Energy’s Office of Biological and Environmental Research (grant no. DE-AC02-5CH11231).

## A Simulation

For a high-resolution phantom, we used a Macrophage cell (jrc macrophage-2) from OpenOrganelle^19^. The Linear Attenuation Coefficient (LAC) values were approximated by measuring the mean gray values from the provided label field. These values were then linearly scaled to match the known LAC values of the respective organelles. The average LAC for the phantom was approximately 0.28 µm^−1^.

We used the linearized incoherent model for the forward model as previously described^10^. The high-resolution and low-resolution PSFs were approximated by modeling the MZP as an ideal circular lens, with the effective PSF calculated according to the methods outlined previously^47^, by the converging illumination emerging from a circular lens aperture based on the Huygens-Fresnel principle^2^.

The MZPs used in the simulations had outer zone widths (OZW) of 35 nm and 60 nm for high and low-resolution setups, respectively. These parameters were chosen to simulate the current configuration at the XM-2 microscope. In XM-2^23^, the condenser is configured as a linear monochromator, formed by the condenser zone plate and a pinhole placed close to the specimen, meaning that we have the same effect of elongated PSF as described in^47^. We have chosen a representative wavelength from the “water window” of 2.4 nm with a monochromaticity of *λ*_0_*/*Δ*λ* = 300. As an “ideal” reference, we used a z-independent PSF equal to the focal point of the 35 nm OZW. For the Beer-Lambert model’s forward and backward projection operators, we used the ASTRA toolbox^34,42,43^ via Tomosipo^20^.

As the phantom’s variability in thickness was high, we used a z-normalizing flat field as the illumination profile so that a single Poisson noise level could better describe the images. This was done by taking an ideal Beer-Lambert projection of the sample and then averaging all illumination profiles over the mean horizontal absorption of the sample.

The approximate noise level of the PSNR was measured from diagonally split experimental data. A similar measurement was done on PSF simulated projections to extract the necessary Poisson count for similar PSNR, with a result of about 150 photons / 10 nm^2^. Code available at https://github.com/ncxt/InteriorSXT.

## B. Experimental results

### B.1. Alignment

The alignment is preceded by the alignment of the full-FOV stack following standard protocol^5^ using AREC3D^36^. Since the entire sample is within the field of view, pre-aligning the capillary using a rigid body transformation is highly robust, even in cases of sparse sampling. The aligned stack is then upscaled to match the resolution of the high-resolution interior projection images.

These images (anchors) ***A***_*i*_ serve as accurately aligned reference images for the interior scan. A schematic representation of the alignment setup is shown in Fig. 8. Note that the mirror images can also be used as anchor points for sampling protocols spanning over 180^°^.

**Figure 8.**
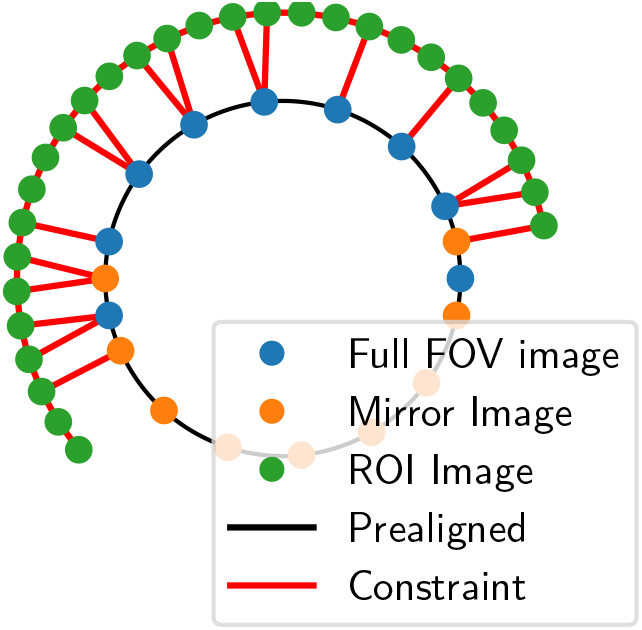
Schematic graph of the alignment procedure. The nodes represent the images, while the edges represent the transformation between them. The inner circle shows the fixed transformation of the pre-alignment, while the outer circle represents the nearest neighbor of the interior scan. An auxiliary constraint is added to a pre-aligned anchor image for an interior image when its angular distance is smaller than that to its nearest neighbor.

We seek translations *T*_*i*_ to the interior images ***I***_*i*_, that maximises the function

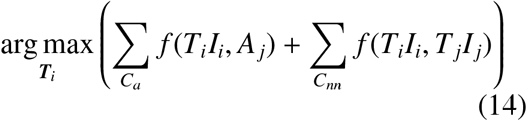

where *C*_*a*_ is the collection of image-anchor pairs *i, j*, for which the angular difference to the anchor is smaller than half of that to its nearest neighbor, and *C*_*nn*_ is the collection of nearest neighbor image-image pairs.

As an alignment metric *f*, we used the normalized cross-correlation

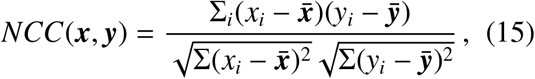

where ***x*** and ***y*** are vector representations of the overlapping information of the two images and 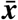, 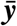, their respective means.

The alignment of the interior images was initially set using a nearest-neighbor approach. A global shift was determined for all images based on reliable full-scan anchor points. This shift was then smoothly interpolated to provide an initial estimate for the interior alignment across the whole dataset. Starting from this initial alignment, the alignment function Eq. (14) was optimized using QuasiNewton methods^44^.

Values of *NCC*(***x, y***) were only evaluated at integer shifts, and intermediate values were obtained by bicubic interpolation. This way, cached values of integer shifts for *NCC* were used to obtain both function values and their derivatives of sub-pixel shifts.

Pure nearest-neighbor alignment generally fares relatively poorly in tomography, as there can be substantial drift in successive alignments. The anchor points (shown in Fig. 8) mitigate this, as it reduces the length of long chains where this drift can happen.

To investigate how many such anchors are experimentally needed for faithful alignment of the interior data, several alignments were performed using only a sparse subset of the complete low-resolution dataset. Fig. 9 presents the mean alignment error as a function of anchor sparsity for experimental bacteria and B-cell datasets. The results demonstrate a slow quality decay (sub-pixel accuracy) with reduced full-FOV images to as low as 10 images, where the alignment error increases significantly.

**Figure 9.**
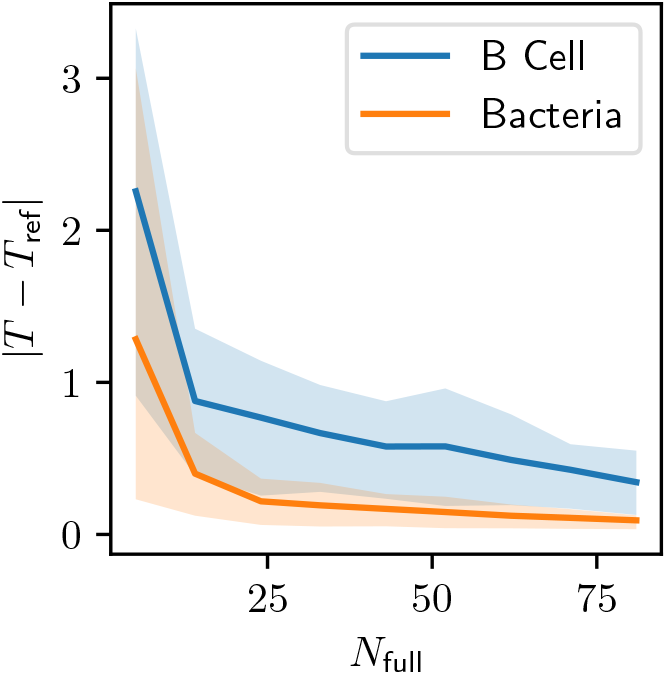
Mean length of the alignment difference between sparse reference data. The shaded area shows the ±*σ* percentile.

Optimal sampling should balance the number of anchor points with alignment stability and dose. Too few anchors lead to significant drift, while excessive anchors introduce sub-optimal dose distribution. Notably, interleaved sampling plays a crucial role in reducing systematic biases by ensuring that mirror images contribute distinct anchor points, thereby enhancing the robustness of the alignment procedure.

### B.2. Note on absorption correction

For both 60 nm and 35 nm OZW objectives, the same monochromator setup is used. In this scenario, a higher-order diffraction of an incoming X-ray beam could be potentially focused on the sample. To account for this potential contamination of X-ray absorption values, we use a dual-energy model to correct the transmission values of the experimental data between sparse and interior scans.

The low-resolution intensity is modeled as pure monochromatic Beer-Lambert

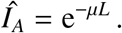

For the high-resolution MZP, we model the intensity as a dual-energy setup, where the intensity is now given by

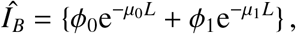

where *ϕ*_0_ and *ϕ*_1_ are the relative intensities of the two energies. As the LAC is energy dependent, we should also describe the sample with two different absorption coefficients *µ*_0_ and *µ*_1_. For the sake of simplicity, we assume that the dependency between absorption and energy can be described as a linear function *µ*_2_ = *cµ*_1_.

With this approximation, the intensity of the high-resolution scan can be described as

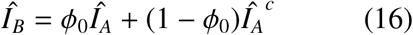

In Fig. 10, we show the best fit of model Eq. (16) to the collected normalized data for the Bacteria sample and compare it to a simple affine model for stray light.

**Figure 10.**
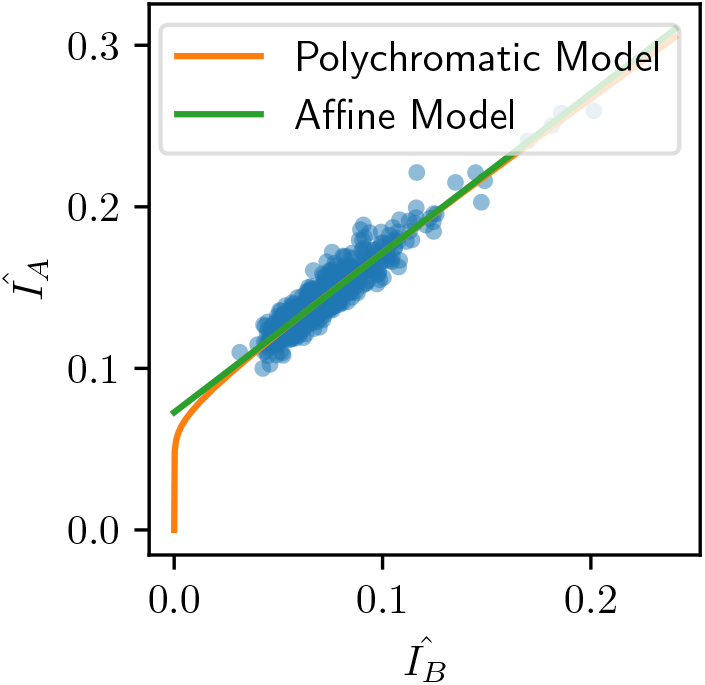
An example of the fitted intensity transform between the 60 nm and the 35 nm images. The orange line shows the used transformation using the polychromatic model of Eq. (16). The green line shows a simple affine model, where the intensity difference is caused by unspecified stray light.

### B.2. Note on bandwidth-limited sampling of the Radon transform

For this, we consider the Radon transform of a two-dimensional impulse function as described by Rattey and Lindgren ^39^. We can express its Radon transformation as

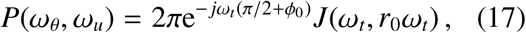

where (*r*_0_, *ϕ*_0_) is the position of the impulse function in polar coordinates, *ω*_*θ*_ and *ω*_*u*_ are the angular and positional frequency, respectively, and the angular frequency *ω*_*θ*_ ∈ ℤ and only takes discrete values.

In a discrete setting where we take the Radon transform for a single voxel of LAC *µ*, this is equivalent to

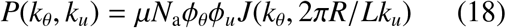

where *ϕ*_*θ*_ and *ϕ*_*u*_ are the appropriate phase shifts for the angular and positional sampling, *N*_*a*_ the number of angles, and *k*_*θ*_ = −*N*_*a*_*/*2 … *N*_*a*_*/*2 and −*L/*2 … *L/*2.

The function |*J*(*x, y*) | approaches zero as *x* ≫ *y*, forming a structure that resembles an “infinite length bowtie”^39^. For the bandwidth-limited case, we multiply the columns by the OTF,

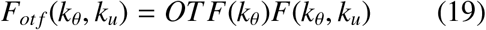

Next, we can investigate the total power collected from different sampling schemes. Integrating the cumulative spectral power of the signal |*F*_*ot f* 2_| ^2^ with respect to *k*_*θ*_, we can quantify the amount of signal power collected by the limited sampling compared to the “ideal case”. In Fig. 11 we an example of the OTF limited Radon transform for the 60 nm OZW and the cumulative power for both 35 nm and 60 nm OZW, showing that most of the signal is collected with sparse sampling while capturing the remaining power requires increasingly dense angular sampling.

**Figure 11.**
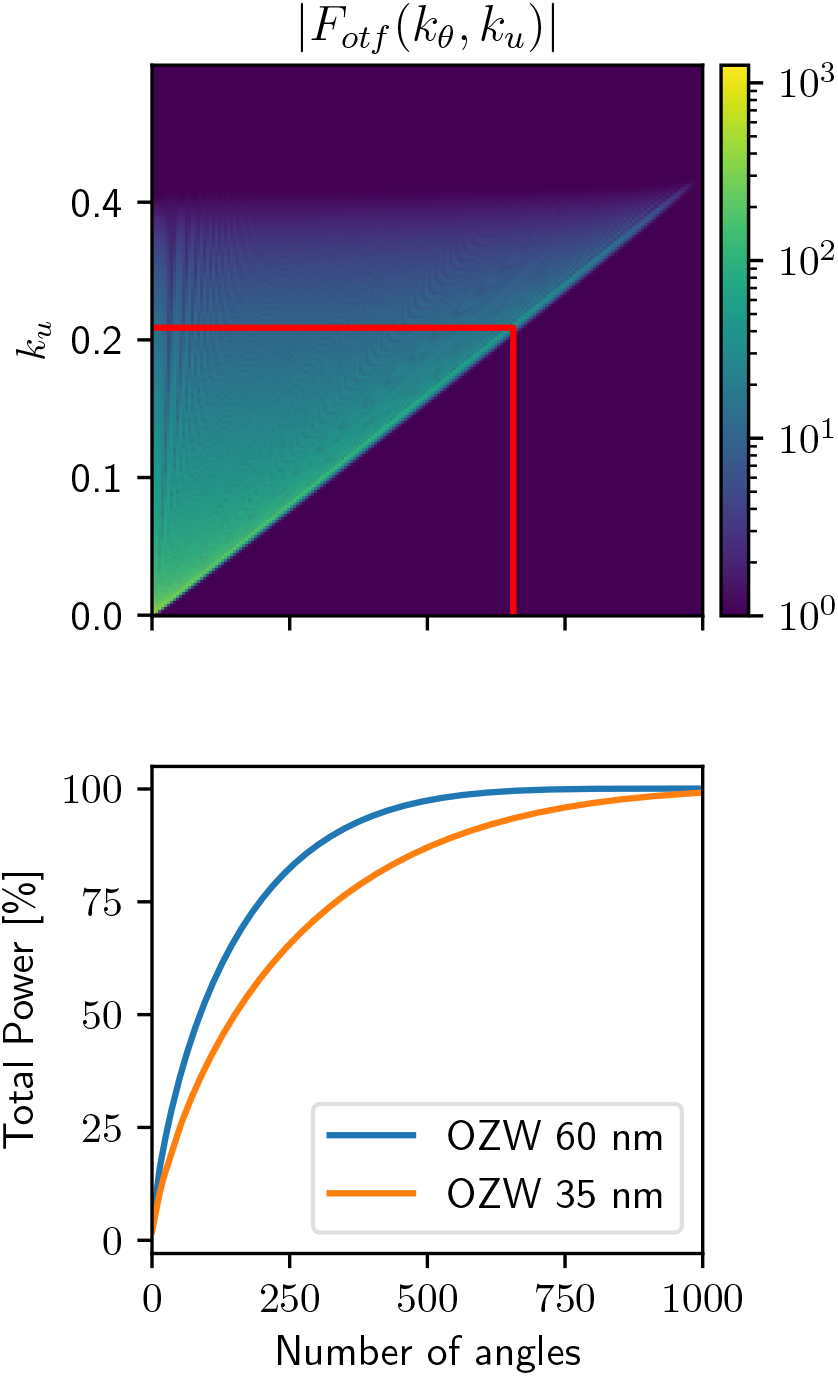
(top) A quadrant of the function |*F*_*ot f*_ (*k*_*θ*_, *k*_*u*_)| showing bandwdth-limited bowtie of the 60 nm OZW. The red line shows the accurate sampling up to 98 % of the OTF power (656 images) (bottom) Integrated total power of the signal with respect to the angular sampling as described in Sec. (B.3). *L* = 800 *r* = 400 sampled with a grid pixel size of 20 nm.

We argue that when the imaging scheme becomes highly noise-limited, the importance of dense angular sampling diminishes, as the SNR of the highest-frequency components degrades. In Fig. 12, we present three examples of sampling with equal total dose. At the lowest dose, no effect of sparse sampling is observed. As the dose increases, the impact of sparse sampling becomes apparent, particularly in cases with the lowest angular sampling densities.

**Figure 12.**
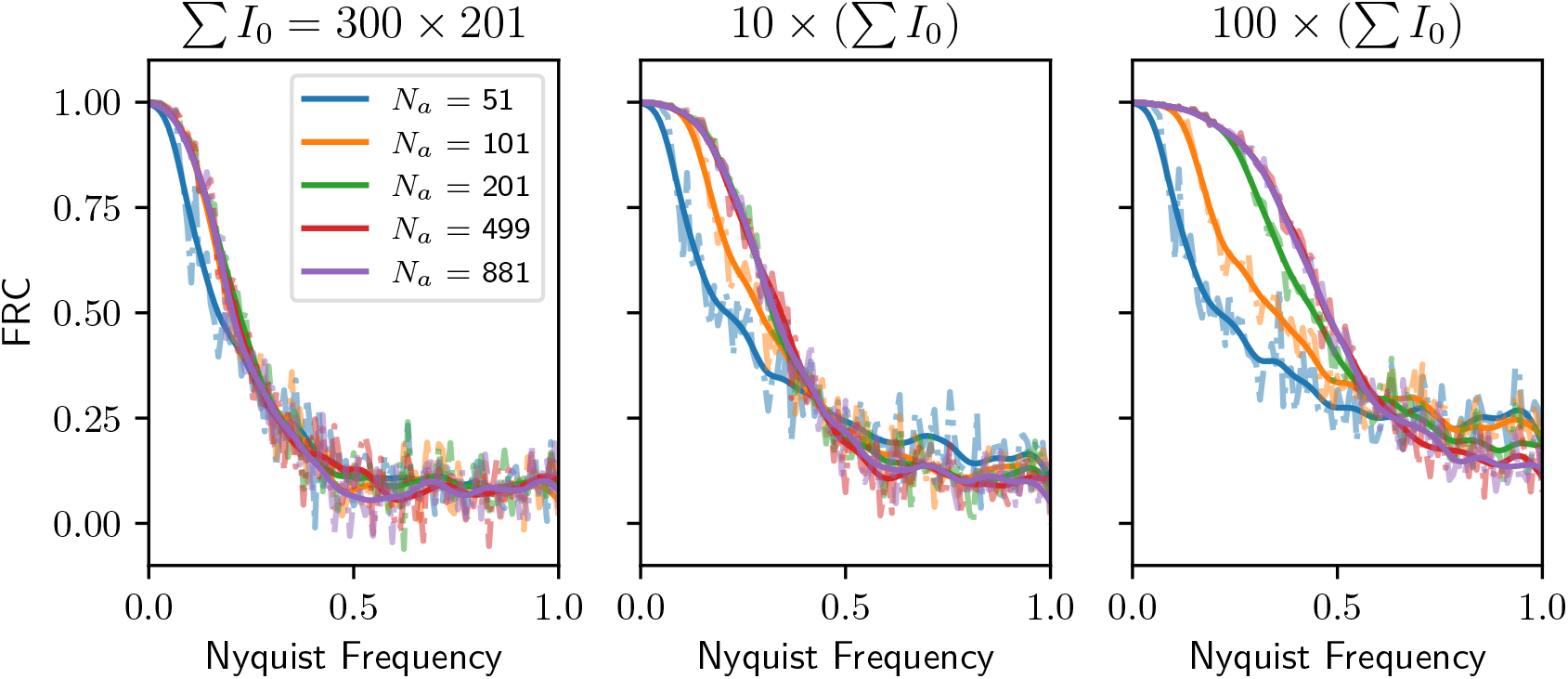
In-slice FRC reconstructions for different angular sampling rates. At lower total doses, dose fractionation holds for sampling rates well below the standard sampling criterion. As the total dose increases, the effects of sparse sampling become more pronounced. However, (a) this occurs at dose levels significantly higher than those typically used and (b) the sampling density in these cases is much sparser than what is employed in practice.

